# Rapid multi-locus sequence typing direct from uncorrected long reads using Krocus

**DOI:** 10.1101/259150

**Authors:** Andrew J. Page, Jacqueline A. Keane

## Abstract

Genome sequencing is rapidly being adopted in reference labs and hospitals for bacterial outbreak investigation and diagnostics where time is critical. Seven gene multi-locus sequence typing is a standard tool for broadly classifying samples into sequence types, allowing, in many cases, to rule a sample in or out of an outbreak, or allowing for general characteristics about a bacterial strain to be inferred. Long read sequencing technologies, such as from PacBio or Oxford Nanopore, can produce read data within minutes of an experiment starting, unlike short read sequencing technologies which require many hours/days. However, the error rates of raw uncorrected long read data are very high. We present *Krocus* which can predict a sequence type directly from uncorrected long reads, and which was designed to consume read data as it is produced, providing results in minutes. It is the only tool which can do this from uncorrected long reads. We tested *Krocus* on over 600 samples sequenced with using long read sequencing technologies from PacBio and Oxford Nanopore. It provides sequence types on average within 90 seconds, with a sensitivity of 94% and specificity of 97%, directly from uncorrected raw sequence reads. The software is written in Python and is available under the open source license GNU GPL version 3.

## Introduction

With rapidly falling costs, long read sequencing technologies, such as from Pacific Biosciences (PacBio) and Oxford Nanopore Technologies (ONT), are beginning to be used for outbreak investigations (Faria et al. 2017; Quick et al. 2015) and for rapid clinical diagnostics (Votintseva et al. 2017). Long read sequencers can produce sequence reads in a matter of minutes compared to short read sequencing technologies which takes hours/days. Seven gene multi-locus sequence typing (MLST) is a widely used classification system for categorising bacteria. It can be used to quickly rule an isolate in or out of an outbreak and knowing a sequence type (ST) can often allow for general characteristics of a bacteria to be inferred. By reducing the time from swab to an actionable answer, genomics can begin to directly influence clinical decisions, with the potential to make a real positive impact for patients (Gardy and Loman 2018).

With the increased speed afforded by long read sequencing technologies comes increased base errors rates. The high error rates inherent in long read sequencing reads require specialised tools to correct the reads (Koren et al. 2017), however these methods have substantial computational resource requirements often taking longer to run than the original time to generate the sequencing data.

A full overview of MLST software for short read sequencing technologies is available in (Page et al. 2017). Of the software reviewed in (Page et al. 2017) only the methods which take a *de novo* assembly as input can be used with long read sequencing technologies, however *de novo* assembly has a substantial post processing computational overhead, which can exceed the time taken to perform the sequencing. StringMLST (Gupta, Jordan, and Rishishwar 2017), was designed to rapidly predict MLST from raw read sets by performing a *k*-mer analysis. MentaLiST (Feijao et al. 2018) takes a similar *k*-mer analysis approach and is designed for large typing schemes, such as cgMLST and wgMLST. They were designed to work only with high base quality short read sequencing data. To our knowledge no method currently exists for calling MLST from uncorrected long read sequencing data.

We present *Krocus* which can rapidly estimate sequence types directly from uncorrected long reads. Results are presented using the largest public dataset of bacterial long read data containing nearly 600 samples generated using the PacBio sequencing technology, and for a small dataset of ONT data. On average it produces sequence correct sequence types in 90 seconds, with a sensitivity of 94% and specificity of 97%. *Krocus* is the only tool which can call MLST directly from uncorrected long reads with high accuracy. It is written completely in Python 3 and is available under the open source licence GNU GPL 3 from https://github.com/andrewjpage/krocus.

## Method

The basic method of *Krocus* is to take short *k*-mers and calculate the coverage over the MLST alleles. As the base errors are mostly uniformly distributed, a well chosen *k*-mer value results in short stretches of error free bases. Some *k*-mers will be erroneously flagged due to errors however as more reads are added (above 5X), these errors are filtered out as they have a low occurrence overall.

*Krocus* takes as input the path to an MLST scheme, a FASTQ file containing uncorrected long reads and a *k*-mer size. The MLST alleles are contained in 7 FASTA files, downloaded from PubMLST (Jolley and Maiden 2010) or taken from the set distributed with the software. Each sequence in the allele files contains a unique identifier and the combination of these allele identifiers gives rise to the sequence type (ST), contained within a profile tab delimited file. An alignment-free *k*-mer sequence analysis approach is used to determine the presence and absence of particular alleles, with optimisations for high error rate long read sequencing data. For a given *k*-mer size, the *k*-mers are extracted from each sequence in each allele file.

In long read sequencing reads, whilst there are high base error rates, the errors are mostly uniformly distributed. The ideal *k*-mer size is the mean number of bases on a read which is free from errors, for example if the base error rate *n* is 91% an error occurs on average every ~11 bases, thus the *k*-mer size *k* calculated as *k* = /i> ⌊*100/(100-n)* ⌋. A *k*-mer size which is too high would invariably always include an erroneous base, reducing the probability of a match with the allele files. A *k*-mer size which is too low would reduce the possible *k*-mer space and lead to an increase in matches by random chance. Each long read is inspected and *k*-mers are generated with a interval of *k* and a step size of *k* giving an average depth of *k*-mer coverage of 1. If a single *k*-mer from this set is present in the allele *k*-mers, the read is kept for further analysis, if no *k*-mers are present, the read is discarded as it is unlikely to contain the MLST genes.

All possible *k*-mers are generated for the read which passed the initial filtering with a step size of 1, giving an average depth of *k*-mer coverage equal to *k,* with k-mers occuring more than 5 times excluded from further comparison as they do not impart useful information. For each allele file, the intersection of the allele *k*-mers and the read *k*-mers is taken. The read is split up into bins of length *k*, and the intersecting *k*-mers are assigned to their corresponding bin in the read to produce an approximate *k*-mer coverage of the read. A sliding window (default 4 times *k*-mer size) is used to span short gaps, which are likely the result of small errors in the underlying sequencing data. The largest contiguous block of *k*-mer coverage in the read is identified, based on the sliding window results, and if it meets the minimum block size (default 150 bases, derived from ⅓ of the average length 467 of all sequences in pubMLST, retrieved 02-02-18), it is said to contain one of the typing alleles. The block is extended by 100 bases on either side to ensure the full allele is captured. The *k*-mers matching this block in the read are extracted and *k*-mer counts corresponding to the allele matching *k*-mers are incremented. The read is reverse complemented and the same search is undertaken once more.

At defined intervals (default 200 reads) the genes of each allele are analysed to calculate the number of *k*-mers covered by the raw read, allowing for the input files to be streamed in real-time as data is generated. If an allele has a gene with 100% *k*-mer coverage, it is said to be present, if it is less than 100%, the allele with the most number of *k*-mers is identified, but with a low confidence flag. Where 2 or more alleles of the same gene have 100% coverage, the sequence with the highest *k*-mer coverage is used. Novel combinations of alleles and new, unseen, alleles cannot be reliably detected using this method, and so are excluded from the analysis.

### PacBio samples

The NCTC 3000 project (http://www.sanger.ac.uk/resources/downloads/bacteria/nctc) aims to produce 3000 bacteria reference sequences using the PacBio long read sequencing technology. Each of the reference strains was selected for sequencing to maximise diversity and to capture historically medically important strains. This is currently the largest public long read sequencing project for bacteria and is still on-going with 1048 assemblies publicly available (accessed 2/1/18). The assemblies were downloaded from the project website and the sequencing reads directly from the European Nucleotide Archive. The sequencing reads were all generated on the PacBio RSII between 2014 and 2017. The assemblies used for comparison with *Krocus* were generated using an open source pipeline (https://github.com/sanger-pathogens/vr-codebase) which first performed a *de novo* assembly using HGAP (SMRT analysis version 2.3.0) (Chin et al. 2013), followed by circularisation with circlator (version 1.5.3) (Hunt et al. 2015), and finally automated polishing with the resequencing protocol (SMRT analysis version 2.3.0) from PacBio.

Each assembly (1048) was sequence typed using the *TS-mlst* software (https://github.com/tseemann/mlst) with ambiguous and untypable samples as identified by *TS-mlst* excluded (339), as a meaningful comparison cannot be made. The *TS-mlst* software was shown in (Page et al. 2017) to never make any false positive ST calls. The dataset was further filtered to exclude samples where there were less than 10 representatives of a species (112) as there were not enough samples to draw any statistically significant conclusions. The remaining 597 samples are detailed in Supplementary Table 1, including accession numbers, and summarized in Table 1, covering 8 species and 537 STs with representatives from both gram positive and gram negative. The FASTQ files of the uncorrected reads were generated from the raw data using the PacBio SMRTlink pipeline (version 5.0.1.9585), and the time for this conversion is not considered in the results presented in this paper as it is a standard preprocessing step required for many downstream analyses. All experiments were performed using the Wellcome Sanger Institute compute infrastructure running Ubuntu 12.04 LTS, with each host containing 32 cores (AMD Opteron Processor 6272) and 256GB of RAM. Only a single core was used in each performance experiment and the mean memory requirement was 0.354GB (std dev 0.16).

**Table 1:** NCTC 3000 PacBio sequenced samples with results after analysis with *Krocus*. An ST is said to be in agreement if it matches the ST called by *TS-mlst* from a *de novo* assembly.

| Species | No. Samples | No. of unique STs | No. in agreement | Mean wall time (s) | Mean Reads |
| --- | --- | --- | --- | --- | --- |
| *E. coli* | 226 | 204 | 204 | 102 | 17524 |
| *E. faecalis* | 11 | 10 | 9 | 42 | 19900 |
| *K. pneumoniae* | 113 | 101 | 108 | 62 | 22297 |
| *P. aeruginosa* | 22 | 21 | 18 | 127 | 41444 |
| *S. aureus* | 114 | 92 | 111 | 122 | 16255 |
| *S. dysgalactiae* | 16 | 16 | 16 | 32 | 10412 |
| *S. enterica* | 48 | 46 | 47 | 107 | 16348 |
| *S. pyogenes* | 47 | 47 | 47 | 37 | 7714 |
| Total | 597 | 537 | 562 | 91 | 17886 |

### PacBio control samples

A set of 74 samples representing 48 species were selected as controls from the NCTC 3000 project. Each were sequenced using the PacBio long read sequencing technology as described previously and are listed in Supplementary Table 2. The controls were selected from within the same genus as the cases as listed in Table 1, but from different species. The species classifications came from experimental techniques. The *de novo* assemblies of each sample were analysed with the *TS-mlst* software, and any which resulted in valid sequence types were removed to reduce the impact of confounders from misclassified isolates.

### ONT samples

To analyse the performance of Krocus on ONT data we use a set of 12 *K. pneumoniae* samples used previously for performance comparisons in (R. Wick, Judd, and Holt 2018) and (R. R. Wick et al. 2017). The dataset is available from (R. Wick 2017a) and detailed in Supplementary Table 3. For comparison the Unicycler assemblies, post Nanopolish, created using only the long read data (R. Wick 2017b) were used.

### Pacbio Results

Each of the assemblies from the NCTC dataset were run through *TS-mlst* to generate a ST. *Krocus* was run for each sample using the uncorrected FASTQ files and default settings and halted when the ST matched the expected result from *TS-mlst.* This replicates the anticipated real-time usage of the software, where a researcher would halt the analysis when a consistent ST result emerges. The time to achieve the correct predicted ST was noted, as were the number of reads, with a mean of 91 seconds, after processing a mean of 17,886 reads. The number of reads required before Krocus correctly predicts the ST is presented in Figure 1a. The running time for each species is presented in Figure 2b. The running time of 3 S. *aureus* samples was elevated due to the need to process a higher than average number of reads, however within 60 seconds 6 out of 7 alleles had been correctly identified, with the last allele taking up to a further 11 minutes to identify correctly. In 94% of cases (sensitivity) the results from *Krocus* and *TS-mlst* were in agreement, with the calculations listed in Supplementary Table 4.

**Figure 1:**
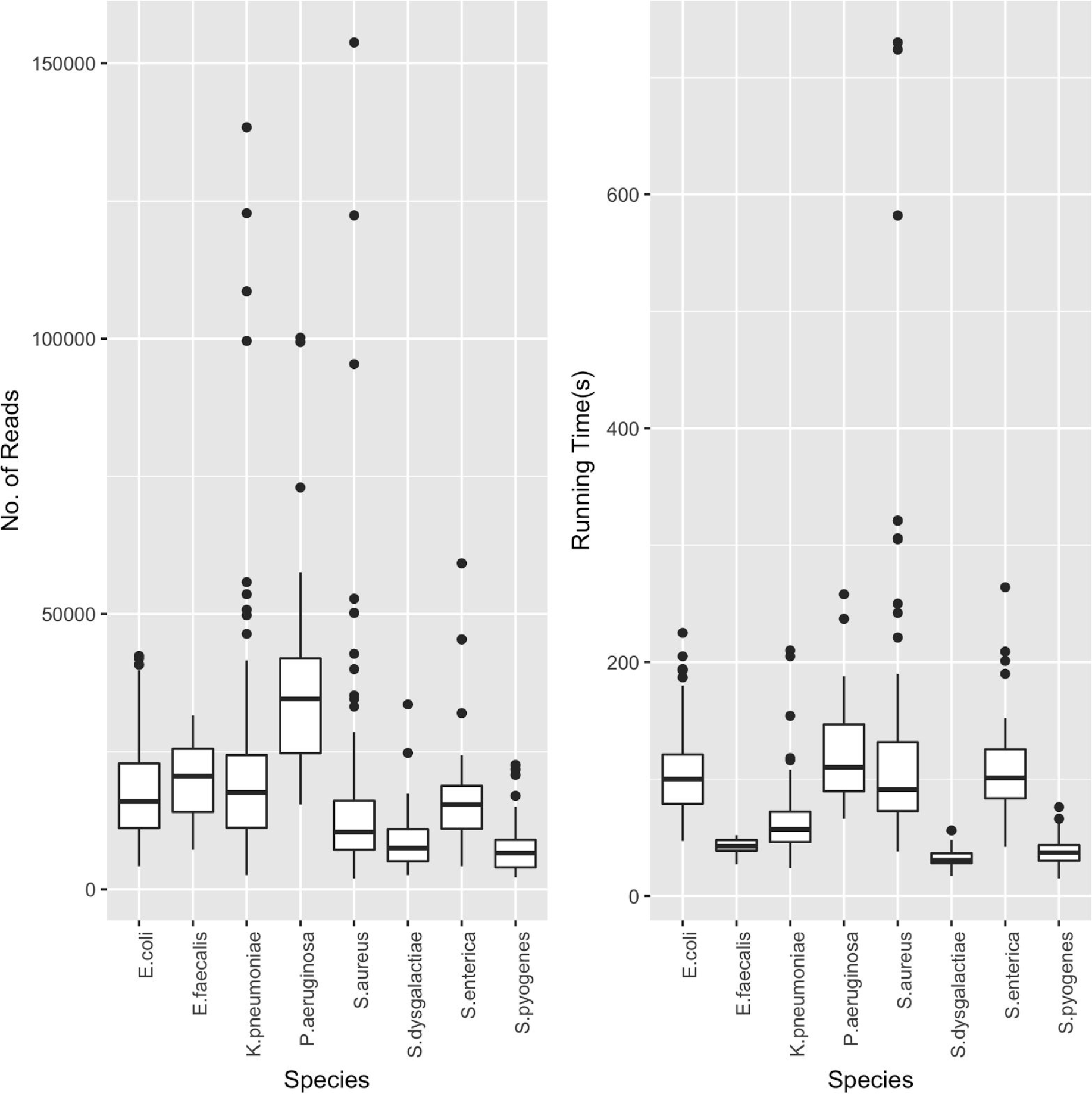
a) Number of reads analysed before the *Krocus* correctly predicted an ST for each PacBio NCTC species analysed, b) time in seconds before *Krocus* correctly predicted an ST for each PacBio NCTC species analysed.

In 35 cases (6%) STs did not match the expected ST or were untypable, with 34 of these calling 6 out of 7 typing genes correctly. In the remaining 1 case 5 out of 7 genes were called correctly. These failures are due to known systematic errors with long homoployers with the PacBio sequencing technology (Quail et al. 2012) which cannot be overcome with short *k*-mers.

The control samples were analysed in a similar fashion to give a specificity of 97.2% (72 out of 74). In the two false negative cases, both contained all 7 typing genes, with one containing 2 copies of gene *phoE* which *Krocus* was unable to distinguish, and one containing a variant in *fumC* which was not in the typing database.

### Nanopore results

For all 12 *K. pneumoniae* samples (100%) of uncorrected ONT reads *Krocus* provided the expected sequence type. The mean time to the expected sequence type was 134 seconds after a mean of 3250 reads. As a comparison, de novo assembled genomes using the ONT reads alone from (R. R. Wick et al. 2017) did not identify any of the sequence types when analysed by *TS-mlst.* This was due to the inherent high base error rate which resulted in a poor quality assembly. Only hybrid assemblies additionally utilising Illumina short read data could be sequence typed. This gives *Krocus* an advantage over *de novo* assembly of ONT only reads.

## Conclusion

*Krocus* is the only tool which can call sequence types directly from uncorrected long reads with high accuracy.The sensitivity of 94% and specificity of 97% achieved by *Krocus* on a large, diverse, PacBio dataset is similar to gold standard experimental standard methods (Liu et al. 2012). By calling sequence types directly from uncorrected long reads, the need for post processing steps and *de novo* assembly is eliminated, reducing the turnaround time for MLST from days to minutes. For a small ONT data, *Krocus* correctly called the sequence type in all cases, compared to *de novo* assemblies of the same data, where no sequence types could be called.

## Acknowledgements

This work was supported by the Wellcome Trust (grant WT 098051). We wish to thank Nick Grayson from the Wellcome Sanger Institute for assistance with the NCTC 3000 dataset. Thanks to João Carriço and Nabil-Fareed Alikhan for providing helpful feedback and suggestions for this paper.

## Supplementary tables

https://docs.google.com/spreadsheets/d/1L6lSp3NJ7PH3alejgq8cn_0Y0aHA0VeFyPYP4yvLpr8/edit?usp=sharing

